# A data-driven approach to automate embolism detection in leaves

**DOI:** 10.64898/2026.07.31.741439

**Authors:** Lorenzo Mandelli, Kate M. Johnson, Stefano Berretti, Maurizio Mencuccini

## Abstract

- Embolism, the formation of air bubbles in the plant water transport system, is a mechanistic driver of plant death. The Optical Vulnerability Technique (OVT) is an imaging method for non-invasive quantification of embolism (including P50, a common metric for drought vulnerability), which can also provide detailed spatial and temporal information. Its major cost lies in the post-processing of thousands of images.
- Here we designed, tested, trained, and make publicly available a neural network model to automate post-processing of OVT images. Using a dataset of 65 leaves from *Senecio pterophorous*, we compared our model predictions to results obtained via traditional post-processing by an expert.
- Our model resolved P50 to within 0.027 MPa of the expert-processed data with training taking 30 minutes to 2.5 hours and model-runtime in the order of seconds to minutes, demonstrating its promise for increasing the efficiency and throughput of P50 calculation. The model’s performance in replicating the pixels that constitute embolism events was lower (mean event-frame IoU of 0.38).
- We invite the community to utilise our model but emphasise that it does not replace the expert-processing pipeline and that care must be taken when considering applying this and similar approaches to OVT data.

## 2 Introduction

Xylem embolism, the rapid formation of air bubbles that blocks the water transport system and leads to tissue damage and death, is the major cause of drought-induced plant mortality globally [IPCC Core Writing Team and (eds.), 2023]. Several approaches can be used to determine plant vulnerability to embolism, from early techniques based on detecting the emission of sounds [Jackson and Grace, 1996], to loss of water-flow capacity [Sperry et al., 1988], [Cochard et al., 1992], [Salleo et al., 1992], [Holbrook et al., 1995], [Pockman et al., 1995] and, more recently, the use of imagery including X-rays [Choat et al., 2015] and optical light [Brodribb et al., 2016a]. In all methods to calculate plant vulnerability to embolism, embolism detection is paired with measurements of plant water stress in controlled experiments to produce ‘vulnerability curves’. A single descriptor often extracted from these vulnerability curves is the P50, the xylem water potential at which half of the water transport system has failed, and this index is used to compare drought vulnerability across different organs, individuals and species.

The Optical Vulnerability Technique (OVT) has emerged as a powerful, non-invasive, image-based method to visualise and quantify plant water transport system failure due to embolism [Brodribb et al., 2016a, Brodribb et al., 2016b]. By using optical light and cameras to observe plant organs, the OVT makes it possible to track embolism continuously, presenting an innovation compared to traditional non-image-based invasive techniques [Cochard et al., 2013], [Gauthey et al., 2020], [Bourbia et al., 2020], particularly in delicate plant organs such as leaves, roots and flowers [Harrison Day et al., 2024], [Johnson et al., 2018], [Bourbia et al., 2020]. Detailed image-based analysis enables not only the obtention of single values, such as P50, for comparison of drought vulnerability, but also highly detailed information on the spatial and temporal spread of embolism. Recent research has used OVT information to determine when and where embolism causes plant tissue death [Tonet et al., 2023], [Brodribb et al., 2021], [Johnson et al., 2022], how the anatomy of the plant water transport system influences the pattern of embolism spread [Johnson et al., 2020]) and how vulnerability to embolism can vary, even within a single organ type, within and across individual plants [Johnson et al., 2022], [Cardoso et al., 2020], [Rodriguez-Dominguez et al., 2018].

The cost of the OVT lies in the image processing. A single experiment can produce hundreds, thousands or even hundreds of thousands of leaf images that must then be post-processed by a trained expert to identify embolism. This workflow entails multiple sequential processing stages and relies heavily on the manual calibration of detection thresholds. A schematic overview of this pipeline is illustrated in Figure 1.

**Figure 1:**
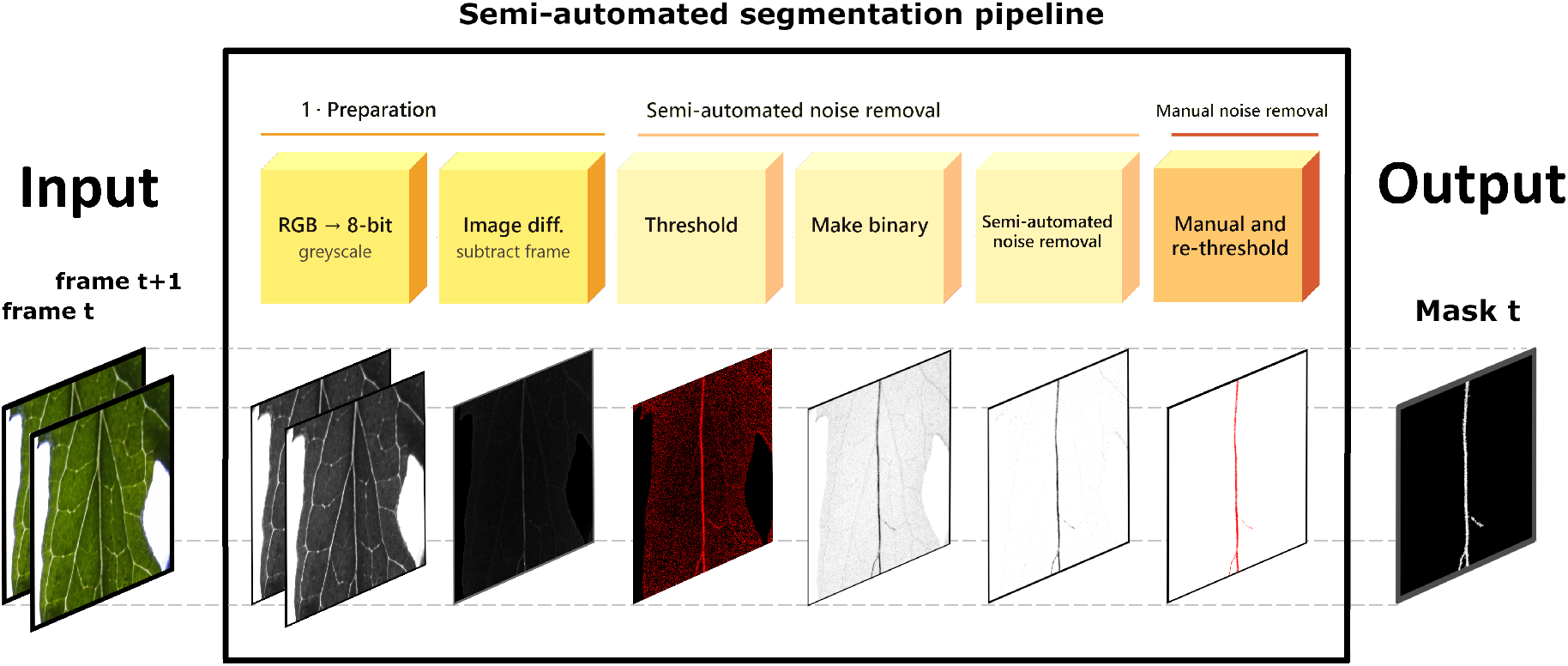
The semi-automated pipeline used by an expert to extract embolism from an OVT image sequence. Starting from the raw frames, which are of a *Senecio pterophorus* leaf in this example, the sequence is converted to greyscale and consecutive frames are subtracted to isolate changes; a signal threshold is then chosen by the user and the result binarized, after which noise is removed in a semi-automated step. A final manual review of the entire processed sequence yields the cumulative embolism mask. The procedure is accurate but labour-intensive and depends on expert judgement at several stages.

The time-consuming nature of the semi-automated pipeline limits the throughput and the breadth of applications of the OVT. Recent research has provided methods to automate the identification of cells in wood [Mei et al., 2026], yet no method currently exists to automate the detection of visual features beyond cell types and in large, temporally sensitive, image sequences, which is particularly relevant in the study of plant water transport. Understanding how and when the plant water transport system fails under stress is essential for predicting drought- and freezing-induced damage and mortality in plants.

Here, we propose a data-driven approach to automate the OVT image-processing pipeline. We design a neural network that, given a pair of consecutive frames, determines whether and where an embolism has occurred in leaves as they dry. Using a dataset of 65 annotated image sequences of *Senecio Pterophorus* leaves we train, validate and test a network that learns to recognise embolism events based on data processed through the semi-automated pipeline. Importantly, we emphasise that our proposed network does not entirely replace the need for semi-automated image-processing conducted by an expert. Rather, we propose that data processed through the semi-automated pipeline can be leveraged to train a model, thereby substantially reducing future reliance on the semi-automated pipeline. This will contribute to increase the throughput of OVT-generated image processing data to enable more efficient quantification of vulnerability to embolism in plants. We also discuss the limitations of our method, emphasise its strengths and weaknesses, suggest how it might be used by the plant science community, and highlight possible next steps for its development.

Our contributions are as follows:

- We design and train a neural network capable of detecting drought-induced P50 in *Senecio pterophorus* leaves with high precision;
- We probe the limits of our approach, investigating both the minimum number of leaves required to train the network to a given performance level to predict P50 and to detect all pixels resolved using the semi-automated pipeline. We also test the ability of our approach to generalise to different species and physical processes (i.e., freeze-thaw embolism) never seen during training;
- We make the code, trained model, and qualitative video results publicly available at https://divanoletto.github.io/Leaf_embolism/.

## 3 Materials and methods

### 3.1 Data Acquisition

The dataset consists of image sequences captured in leaves as they dried, which were post-processed by an expert to identify air embolisms in the venation networks. Data were obtained from 3-5 leaves per plant in 15 plants of a single species, *Senecio Pterophorus*. Each leaf was monitored with a specialised imaging clamp [Cavicam, 2026]. These custom clamps comprise a raspberry pi single-board computer, raspberry pi camera, and customised software designed to allow continuous image capture at a fixed magnification under LED illumination. The images were acquired automatically every 5 minutes, producing image sequences that track individual leaves from a well-hydrated state to complete desiccation (from 0 to 100% embolism). Plant water potential was simultaneously measured on the main stem of each plant during drying using a Stem Psychrometer (ICT PSY, Armidale, NSW, Australia) following [Johnson et al., 2022]. The cavicams are a streamlined method for applying the Optical Vulnerability Technique (OVT) [Brodribb et al., 2016a], the use of optical light and cameras to visualise and quantify air embolism.

Post-processing of the images was conducted using the well-established OVT semi-automated image-processing methodology outlined in [OpenSourceOV, 2022], and in numerous publications [Brodribb et al., 2021, Johnson et al., 2018, Bourbia et al., 2020, Cardoso et al., 2020, Brodribb et al., 2016a, Brodribb et al., 2017]. In brief, the rapid change in light transmission that occurs when water-filled veins rapidly fill with air (embolise) can be detected visually. Semi-automated post-processing consists of conducting an *image difference*, where each image is subtracted from the one before, to reveal the changes between them, followed by various steps to isolate the rapid embolism events and distinguish them from other visual changes, such as sample movement and shrinking, that occur as a sample dries (see Figure 2). While this dataset contains leaves of a single species, experiments to test the generalisability of the proposed method on new species and unseen conditions can be found in Section 4.4.

**Figure 2:**
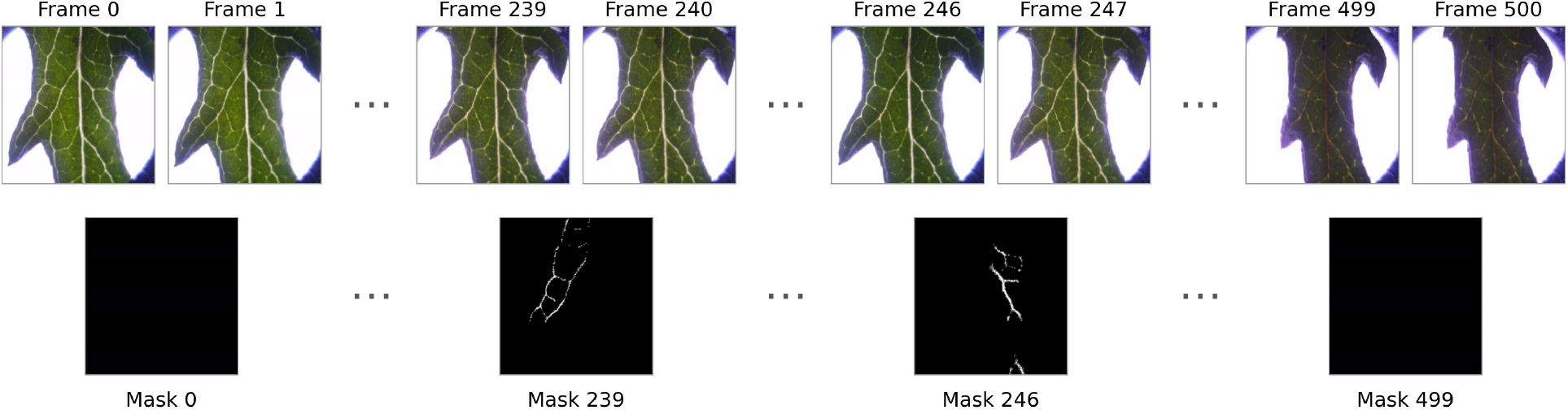
Example sequence from the dataset for a single leaf of *Senecio pterophorus*. The top row shows the optical frames acquired over the course of the drought experiment, while the bottom row shows the corresponding embolism masks, in which bright pixels mark the vein segments that embolise between consecutive frames. Sequences of three dots denote the many intermediate frames omitted for clarity. As dehydration progresses, the leaf undergoes pronounced morphological change: the lamina steadily shrinks, curls and contorts, so that its silhouette at the end of the sequence bears little resemblance to its fully-hydrated shape at the start.

Embolism events were annotated as binary ground-truth masks: for each pair of consecutive frames, a mask marks the pixels where a new air-filled (embolised) region appears between the earlier and the later frame. These masks are the reference against which the model is trained and evaluated. Because the network operates on consecutive frame pairs, a sequence of *N* image frames yields *N* − 1 frame pairs (and the corresponding *N* − 1 masks).

### 3.2 Dataset Composition and Splits

The dataset comprised 65 leaf sequences, for a total of 30,367 individual frames. The sequences were partitioned at the *leaf level* into three disjoint sets (training, validation, and test) so that no leaf appeared in more than one set. Splitting at the leaf level (rather than mixing frames from the same leaf across sets) is essential: consecutive frames of a single leaf are highly similar, and allowing them to appear in both training and test would let the model effectively *memorise* a leaf and produce over-optimistic results. The split sizes are summarised in Table S1.

From a deep learning perspective, a peculiar and challenging property of this dataset is the fact that embolism only occurs in a small portion of the images, both temporally across the sequence, and within the area of each frame. Only about 9% of all frame pairs contained any embolism event, and even within those pairs each embolism event usually covered only a small fraction of the image, the rest being unchanged background. A naive model could therefore reach very high pixel accuracy simply by predicting *no embolism* everywhere, rendering it useless in practice. Addressing this challenge is the main reason behind the choices of the loss function and sampling strategy described below.

Our goal was to create an automatic, data-driven method that, following training on data processed through the semi-automated pipeline, could be given a sequence of images of leaves subjected to a stress factor (e.g., drought) and quickly identified which vessels underwent embolism at each frame. To this end, we trained a neural network on data annotated by an expert, so that, given a pair of consecutive images (frame *t* and frame *t* + 1), the network could predict whether and in which pixels an embolism had occurred. By applying this prediction to every pair of frames, we could obtain a vulnerability curve, the P50 and an overall map of the embolised vein network. Figure 3 summarises the proposed pipeline.

**Figure 3:**
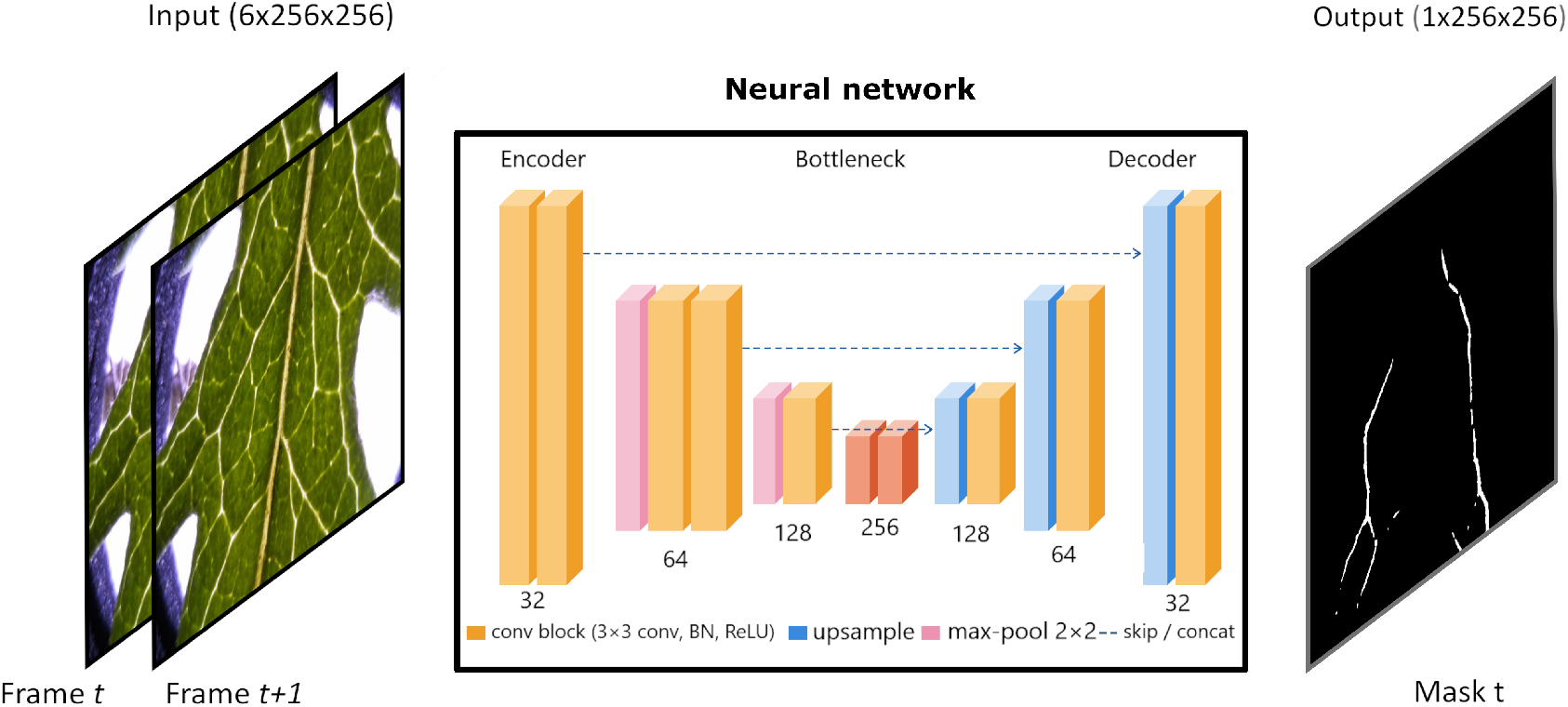
Overview of the proposed neural network approach applied to a *Senecio pterophorus* leaf sequence. Two consecutive frames (frame *t* and frame *t* + 1) are fed into a U-Net-style neural network, which outputs a mask for frame *t* indicating the pixels where embolism has occurred.

In the following sections, we describe the architectural characteristics of the neural network we use (Section 3.3), the loss functions implemented to guide the network toward learning meaningful information (Section 3.4), and the data augmentation and over-sampling methods adopted to focus training on the rare and small regions of the image where embolisms occur (Section 3.5).

### 3.3 Network Architecture

We used a convolutional neural network with a U-Net architecture [Ronneberger et al., 2015], a design that has become a standard for biomedical and microscopy image segmentation. A U-Net works in two stages: an *encoder* progressively compresses the input image into a compact representation that captures which features are present and at what scale, and a *decoder* reconstructs a full-resolution output from it, recovering where each feature is located. *Skip connections* [He et al., 2016] pass spatial detail directly from the encoder to the decoder, so that fine structures are not lost in the compression. A scheme of the network architecture is shown in Figure 3. The key design choice for our task is the **input**: the two consecutive RGB frames are stacked together into a single six-channel input. Importantly, the network considers frame-pairs with no knowledge of which sequences they belong to or their temporal position in the sequences. Presenting frames-pairs lets the network compare them internally and respond to the *changes* between them, rather than to the static appearance of either frame alone. The **output** is a probability map at the same resolution as the input, in which each pixel value is the model’s estimated probability that a new embolism event has occurred there. This map was thresholded at 0.5 to obtain the final binary embolism mask. To limit overfitting on a training set of limited size, the network also used dropout [Srivastava et al., 2014] during training, to improve generalisation to unseen leaves.

### 3.4 Loss Function

The network was trained by minimising a loss function designed to cope with the small number of frames and small areas within frames that are occupied by embolism events. A standard pixel-wise loss would have been dominated by the abundant background and would have barely been affected by the few pixels of interest that constitute the embolism events. We combined, with equal weight, two complementary terms. The *Focal loss* [Lin et al., 2018] down-weights pixels that are already classified correctly with high confidence, so training concentrates on the scarce, difficult embolism pixels rather than on the background. The *Dice loss* [Milletari et al., 2016] takes a region-level view, measuring the overlap between the predicted and the true embolised region as a whole; because it depends only on this overlap, it stays informative even when the embolised area is very small. Used together, the two terms make the model focus on the rare embolism events both at the level of individual pixels and at the level of whole regions.

### 3.5 Patch Sampling and Data Augmentation

For computational reasons each frame was resized to 512 × 512 pixels, a resolution at which embolism events remain clearly visible, and the network was trained on smaller 256 × 256 crops rather than on whole frames. Because frame pairs containing an embolism event are rare, they were *oversampled* during training and the crop was centred on the embolised region (with a small random offset), so that the model was shown the events of interest far more often than their natural frequency would allow. To increase the variability of the training set, each cropped area was also randomly transformed with geometric and photometric augmentations (flips, rotations, mild elastic deformations, and small changes in brightness, gamma and sensor noise). These transformations were chosen to mimic realistic differences between cameras, sessions and consecutive readouts, but also the leaf movement and shrinkage observed during each sequence (Figure 2), so the augmented images remained physically plausible.

### 3.6 Evaluation Metrics

We assessed the model along two complementary axes: how faithfully it reproduced the progression of embolism over time including the estimation of the P50 (leaf-level quality) and how accurately it delineated each embolism event identified in the ground-truth data (pixel-level segmentation quality).

#### FP50 and P50

To quantify temporal accuracy, we accumulated the embolised area frame by frame and recorded *FP50*, the *frame index* at which 50% of the total embolised area of a sequence was reached. We employed FP50 in two ways. The first case was against the *water potential* : Since plant water potential was measured throughout each experiment, we could match the timing of FP50 with the timing of the measured potentials, obtaining a physiologically meaningful descriptor of the leaf’s vulnerability to embolism, the P50. We then compared model-predicted P50 against ground-truth P50 (i.e., P50 derived from expert processing through the semi-automated pipeline). The second case did not instead consider the actual water potentials (for the experiments reported in the Supplementary Information). Here, we employed FP50 directly (i.e., the *frame index*), measuring whether the predicted amount of embolism was placed correctly in time by the model, and then compared predicted versus observed *frame index*, to reveal whether the model recovered the *timing* of embolism, independently of small pixel-level errors. An example of the plot of %embolism versus water potential for one leaf is shown in Figure 4(a).

**Figure 4:**
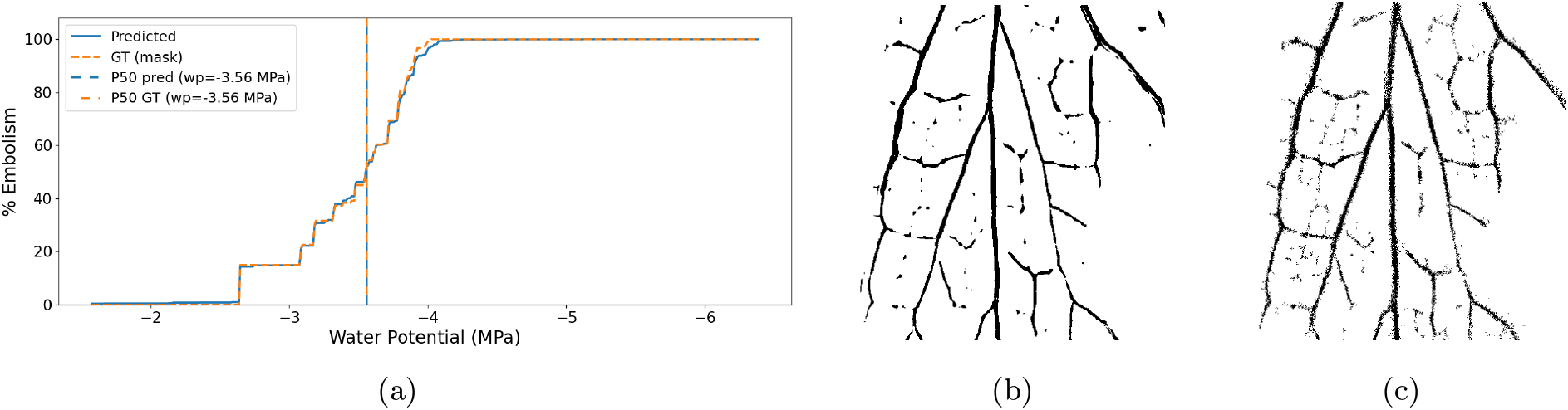
Qualitative evaluation of the network results on *Senecio pterophoru*s from the test dataset. **(a)** Comparison of embolism accumulation over time between model predictions and ground truth (derived from expert image processing using the semi-automated pipeline), along with the corresponding predicted and ground-truth P50 values; **(b)** predicted cumulative mask; **(c)** corresponding ground truth cumulative mask. Legend (left-hand panel): the solid blue line (“Predicted”) shows the change in %embolism as a function of water potential predicted by the U-Net model; the dashed orange line (“GT (mask)”) shows the change in %embolism as a function of water potential in the annotated images (ground-truth); the vertical dotted blue line marks the P50 predicted by the U-Net model; and the vertical dotted orange line marks the P50 obtained from manual image annotation.

#### Cumulative mask

For a qualitative, whole-sequence comparison, we overlaid all per-frame predictions of a sequence into a single *cumulative mask*, and likewise for the ground truth. Placing the two side by side gives an immediate visual impression of whether the model recovers the overall spatial pattern of embolism across the leaf, independently of the exact frame in which each embolism was detected. An example is given in Figure 4, where the predicted (b) and ground truth (c) cumulative masks are shown.

#### Segmentation quality (IoU)

At the pixel level we summarised how well the predicted and the ground-truth masks overlapped with the *Intersection-over-Union* (IoU) [Everingham et al., 2010], a standard segmentation score ranging from 0 (no overlap) to 1 (perfect overlap). Because the vast majority of frame pairs contain no embolism at all, where the IoU is not meaningful, we report it averaged over *event frames* only (the frame pairs whose ground truth contains at least one embolised pixel), which is where segmentation quality can actually be measured.

#### Precision and Recall

While the IoU summarises overall segmentation quality in a single number, it does not reveal which are the *type of mistakes* the model makes. For this purpose, we additionally report precision and recall, again restricted to event frames. *Precision* is the fraction of the pixels flagged as embolism that are actually embolised, so a low precision indicates over-segmentation (the model marks embolism where there is none). *Recall* is the fraction of the truly embolised pixels that the model recovers, so a low recall indicates under-segmentation (the model misses part of the embolised area). Reporting the two separately distinguishes these opposite failure modes.

## 4 Results

### 4.1 Testset Results

Across the ten test leaves (testset), the predicted P50 matched the ground truth to within 0.027 MPa on average (5a). This is also visible in Figure 5b, where the predicted and ground-truth P50 fall at nearly the same water potential. The close match between the data processed through the semi-automated pipeline and model-predicted data can also be seen qualitatively (Figure S3). At the pixel level, the model reached a mean event-frame IoU of 0.38, with similar precision and recall values of 0.56 and 0.55, respectively (5a).

**Figure 5:**
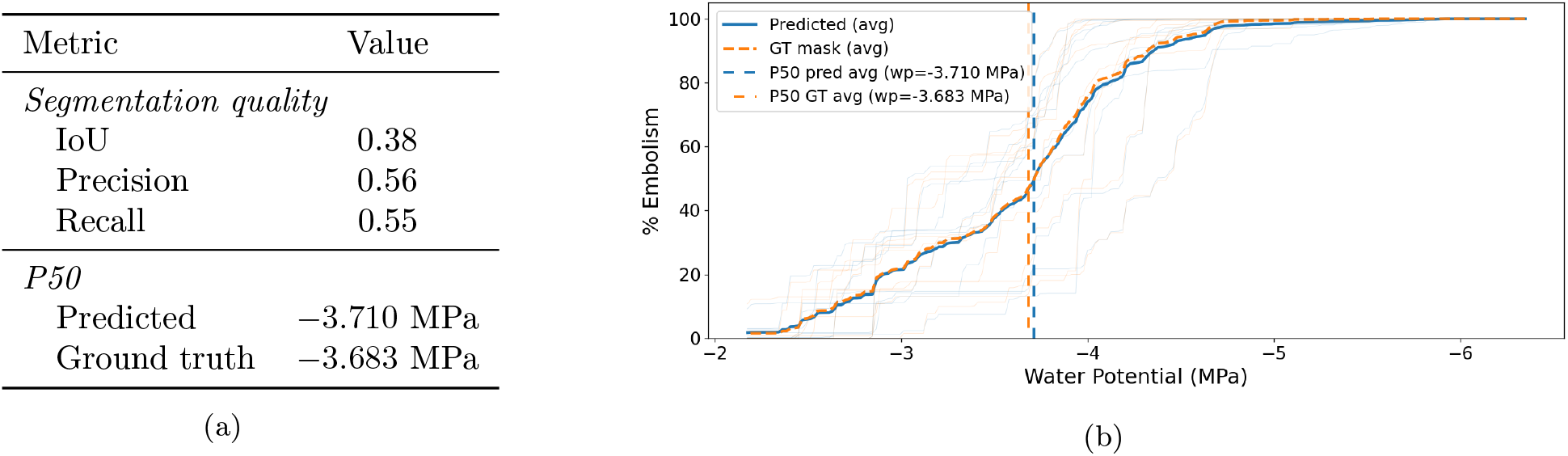
Results on the ten test leaves of *Senecio pterophorus*. **(a)** Mean segmentation metrics (event frames) together with the mean P50, model-predicted vs. ground truth (derived from expert image processing using the semi-automated pipeline). **(b)** Percentage of total embolism, predicted vs. ground truth, as a function of leaf water potential, averaged across the test sequences. Legend (left-hand panel): the solid blue line (“Predicted”) shows the change in %embolism as a function of water potential predicted by the U-Net model; the dashed orange line (“GT (mask)”) shows the change in %embolism as a function of water potential in the annotated images (ground-truth); the vertical dotted blue line marks the P50 predicted by the U-Net model; and the vertical dotted orange line marks the P50 obtained from manual image annotation.

### 4.2 How Many Leaves Are Needed for training?

A practical question for adopting this approach is how much annotated data is required to achieve accurate predictions of P50 and high resolution of embolised pixels. To answer this question, we trained the model on random subsets of the training set, incrementally increasing the number of leaves *N* for sampling, where *N* ∈ {4, 8, 12, 16, 24, 32, 40, 46}. For each size, we repeated the experiment with several independent random draws of the subset (from ten draws at *N* = 4 down to a single draw at the full *N* = 46) and report the mean and standard deviation across draws of the difference between the ground truth and the predicted P50 and the IoU metrics, so that the learning curves reflect not only the average performance but also the model’s sensitivity to *which* leaves happen to be in the training set (Figure 6).

**Figure 6:**
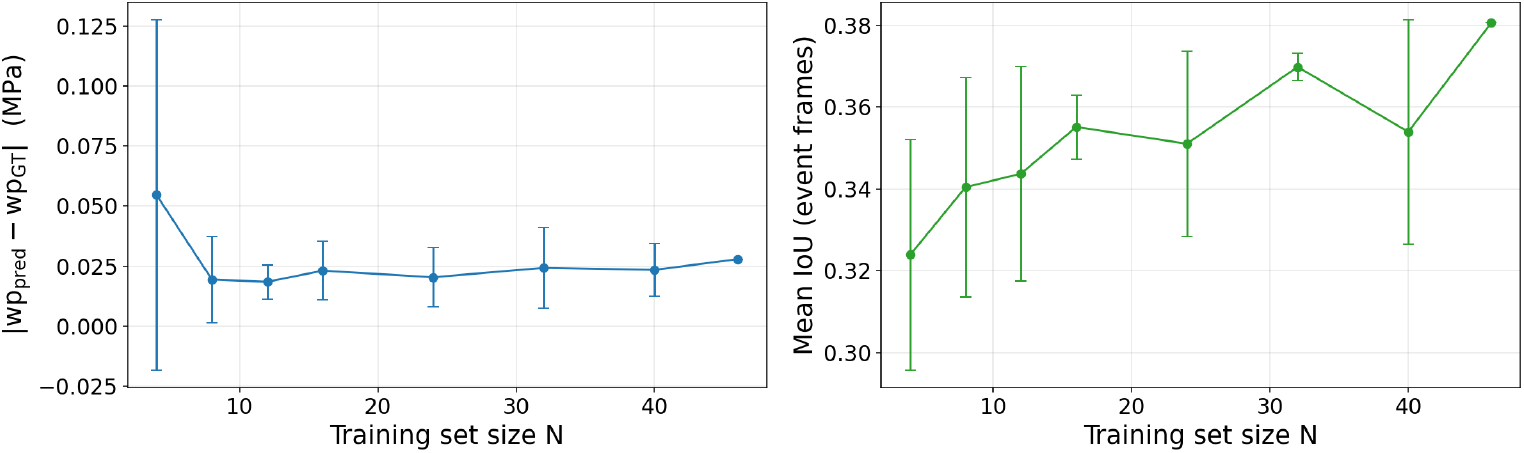
Learning curves as a function of the training set size *N* (number of training leaves from the *Senecio pterophorus* dataset). **Left:** absolute error (i.e., difference) between the predicted and ground-truth leaf water potential at %50 embolism (lower is better). **Right:** mean IoU (Interception over Union) on event frames (higher is better). Both metrics improve rapidly with the first few training leaves. For the absolute error, a plateau is observed: at around 0.02 MPa just using *N* ≈ 8 leaves; the segmentation IoU, instead, continues to improve with more data, even if with different velocity, reaching its best value when the full training set is used. The vertical bars at each point give the standard deviation at each sampling intensity.

The error in predicted P50 dropped sharply over the first few leaves, then a plateau was observed with as few as *N* ≈ 8 leaves at which point the model estimated the water potential at 50% embolism to within roughly 0.02 MPa, and adding more leaves did not result in a further improvement. In contrast, the event-frame IoU kept improving gradually with more data and reached its best value only at the full training set, showing that fine pixel-level mask quality, unlike the aggregated P50 readout, continued to benefit from additional leaves even after the improvement in P50 prediction had saturated.

### 4.3 How much time does it take?

In all our experiments, we used an NVIDIA GeForce RTX 5090 GPU with a training batch size of 64 and an inference batch size of 1 (where batch size represents the number of images processed simultaneously in a single forward pass). Training the model on the total training set of 46 leaves took 2.5 hours; to train the model on the 8 leaves needed for P50 estimation to within 0.02 MPa (sec 4.2) took 30 minutes. Once trained, inference on a 500-image sequence takes around 7.5 seconds on the GPU and around 2.5 minutes on the CPU alone (AMD Ryzen 7 9800X3D), the time needed scaling proportionally with sequence length (around 15 seconds for 1000 images on the GPU). Inference is therefore feasible in seconds to a few minutes even without dedicated GPU hardware. Conversely, processing of a single 500-image sequence using the semi-automated processing pipeline (Figure 1) can take anywhere from 20 minutes to several hours. Considering an intermediate estimate of 1 hour per image sequence for the semi-automated pipeline, with 8 leaves needed for accurate P50 estimation, the total time needed (manually processing sequences, train the model, run the model on a new 500-image sequence) is 8 h 30 min in total. Crucially, this is a one-off cost for a model trained on a singular organ of a single species: once trained, the model processes each additional 500-image sequence in seconds to minutes rather than roughly an hour, making the approach increasingly advantageous as the number of sequences grows.

### 4.4 How well does the model generalise to unseen scenarios?

We conducted two tests to explore the ability of the model to generalise beyond the conditions under which it was trained. We tested the model by applying it without any retraining or fine-tuning (zero-shot), to two leaves of a different species (i.e., *Liriodendron tulipifera*), one which had been exposed to drought stress and the other exposed to freezing and thawing. In neither of these cases was water potential (MPa) measured, so only the FP50 was estimated, not the P50. In the drought-stressed *L.tulipifera* leaf, shown in Figure S1, the spatial patterning of embolism is well resolved, with a similar event frame IoU of 0.34 to that found in *S. pterophorus*, i.e., 0.38 and a particularly high precision (0.69), meaning the pixels it flags as embolism are mostly correct, though its recall is lower (0.39), so it misses part of the embolised area. This is also apparent in the cumulative masks (Figure S1(c),(d)). The predicted and ground-truth patterns share the same overall layout, but the prediction is visibly sparser, with parts of the embolised network omitted (Figure S1). There is a large mismatch in the predicted (frame 190) and ground truth FP50 (frame 106) (Figure S1(a)). The segmentation quality in the freeze-thaw leaf is substantially lower (event-frame IoU 0.12) compared to the drought examples (Figure S2). However, there is a near perfect match between the predicted FP50 frame (162) and ground truth (163).

## 5 Discussion

Here, we develop a data-*driven* approach for the detection of air embolism in image-sequences of leaves. We present a model that can be trained using data from an existing and widely used semi-automated image-processing pipeline, to allow further data to be processed through a single neural network. Given a pair of consecutive frames, the neural network predicts whether and where a new embolism event has occurred. Aggregating these predictions over a whole image sequence from a single leaf yields a vulnerability curve, from which P50 can be extracted, along with a cumulative embolism map.

Our experiments show that, despite the modelling challenges presented by the relatively small proportion of the sequences and image areas occupied by embolism, the model recovers the physiologically meaningful quantity, P50, with high fidelity. The close match between the predicted P50 and that derived through the semi-automated processing pipeline (within 0.027 MPa on average) demonstrates that this model is a promising tool to increase the efficiency of quantifying plant vulnerability to embolism.

The exact match in pixels between the groundtruth and predicted embolism masks is relatively low (event frame IoU: 0.38), but this value must be interpreted in light of the nature of the reference data rather than read as a straightforward measure of model failure. Ground-truth masks derived from OVT image processing are obtained by thresholding frame-to-frame differences, and are therefore intrinsically speckled (4c): within a single embolism event, individual pixels are labelled independently, so that the annotated region is a fragmented, non-contiguous cluster of positive pixels interleaved with negative ones. The neural network, by contrast, predicts spatially continuous regions effectively identifying that an embolism has occurred within a small area rather than reproducing the exact pixel-level texture of the reference (Figures 4b, S3). Because IoU is computed pixel-wise, and because embolism events are thin and sparse, the IoU is is heavily penalised. An IoU of 0.38 in structures of this kind is therefore not comparable to the same value in conventional segmentation benchmarks.

The fact that the mismatch in pixels between the ground-truth and the model is largely textural rather than locational is confirmed by the cumulative masks in Figures 4 and S3: while the pixel-level pattern of each individual event differs from the reference, most of the embolised area is resolved faithfully. The same conclusion follows from the similarity between precision and recall, which indicates that the model neither systematically over-segments (which would depress precision) nor under-segments (which would depress recall), but makes errors of both kinds in roughly equal measure. The per-frame masks are thus imperfect in their fine detail, as expected for such thin, sparse structures, but they are not systematically biased.

The diffuse nature of the embolism events in the higher-order (smaller) veins means that these are the ones most often lost in the model predictions (Figure S3). As the brightness change is fainter and covers fewer pixels, the network is correspondingly less confident, and the weakest events fall below the threshold and are dropped. The continuous improvement in pixel-level segmentation quality with the addition of more leaves (Figure 6), with no sign of plateau, indicates that the finest structures are not yet fully ‘learned’ at the annotation volumes used here. This does not, however, propagate to the vulnerability curve. Embolism in the smallest veins accounts for only a small fraction of the total embolised area, and the aggregation involved in computing the curve averages out local, pixel-level errors. Accordingly, the P50 readout plateaus with as few as eight training leaves, while segmentation quality is still improving: a reliable P50 can be recovered with a comparatively modest annotation effort, even from a network that does not segment perfectly.

While diffuse embolism and embolism in the smaller veins constitute a small proportion of the total percentage, meaning that omission or partial resolution of these by the model did not have a large effect on the vulnerability curve or the P50, losing these would have negative consequences in studies where these details are critical. This includes research focusing on the spatial or temporal pattern of embolism spread [Johnson et al., 2020], vein structure, path-length, and vein order ([Lu et al., 2025], [Tonet et al., 2023]). The model presented here and similar approaches may not be appropriate in these cases or may require significantly more leaves for training. One of the strengths of non-invasive imaging techniques such as the OVT is the rich and detailed spatio-temporal information that can be harnessed. This detail of course, can and often is, lost in the semi-automated processing step as well. We therefore suggest that individual researchers critically assess the goals of their study before deciding how to conduct post-processing.

We also wish to comment on the generalisability of our model beyond the settings evaluated here. Our model was trained and tested on a single species, with ground truth defined by a single expert, and agreement is expected to be highest under exactly these conditions: the network learns not only the visual signature of embolism, but also the particular conventions and thresholds applied by that annotator. These two limitations are distinct and call for different remedies. *Annotator dependence* could be reduced by training on ground truth pooled from multiple experts: the network would then converge on the consensus interpretation rather than reproducing any single set of conventions. This may lower apparent agreement with any individual annotator while making the output less biased. *Species dependence*, by contrast, is a matter of domain rather than convention, and may be better addressed by training a general model on a dataset spanning multiple species and imaging conditions. Conversely, there is also an opportunity to create models specific to species or organs to maximise the accuracy of embolism detection at the species or organ level in plants.

Information on what a more general model, or specialised models for species or organs, would need to accommodate can be derived from applying our model to unseen circumstances. Our results suggest that the network captures a genuinely transferable visual signature of embolism, but also make clear the limits of applying it outside its training domain. While much of the embolism in the *L. tulipifera* leaf exposed to drought was detected by the model, the omission of some embolism, including the more diffuse embolism in the major vein created temporal inaccuracy in the FP50 prediction (Figure S1). In the *L. tulipifera* leaf exposed to freezing and thawing, only the earliest freeze-thaw embolism events were detected along with the signal of thawing in some of the leaf mesophyll, yet the FP50 was resolved with high fidelity. The match in FP50 can be explained by the rapid occurrence of embolism in this case, occurring only in the largest veins and rapidly over three images (15 minutes) in contrast to drought-induced embolism, which occurs in all vein orders and over hundreds of images (hours to days) [Johnson et al., 2025]. The close match between the FP50s despite the very low match in embolism resolved is a reminder of the importance of consulting the visual information produced by the OVT, rather than relying solely on the numerical output. Additionally, like *S.pterophorous, L. tulipifera* possesses reticulate venation, so in some sense it can be considered an ‘easy’ test.

Our results suggest that a data-driven approach can substantially increase the throughput of OVT analysis for the calculation of P50, broadening the practical applicability of the technique. Several directions remain open for future work. Extending the model to venation patterns, species, expert conventions, and stress factors beyond those examined here will require broadening the training set accordingly. We hope that the public release of the code and the trained model will encourage further work on automating embolism analysis and on benchmarking against expert-processed data.

## 6 Acknowledgements

KMJ was funded by the European Union on a Marie Skłodowska-Curie Actions (MSCA) Postdoctoral fellowship, 101107177—IVERdrought. Views and opinions expressed are, however, those of the author(s) only and do not necessarily reflect those of the European Union or the MSCA. Neither the European Union nor the granting authority can be held responsible for them. MM acknowledges support from the Spanish Ministry of Science and Innovation (MICINN) through grant PID2022-137270NB-I00 (DRASTIC) and from the EU Horizon 2020 programme through grant 862221 (FORGENIUS).

## 8 Author contributions

MM, LM and KMJ conceived and developed the idea. KMJ collected and processed the training dataset. LM designed, ran and reported on the model and wrote the first draft of the manuscript. LM, KMJ and MM interpreted the results and wrote the final manuscript with contributions from SB.

## 9 Data Availability

The model and code are available at this link: https://github.com/divanoLetto/.

## Supporting Information

**Table S1:**
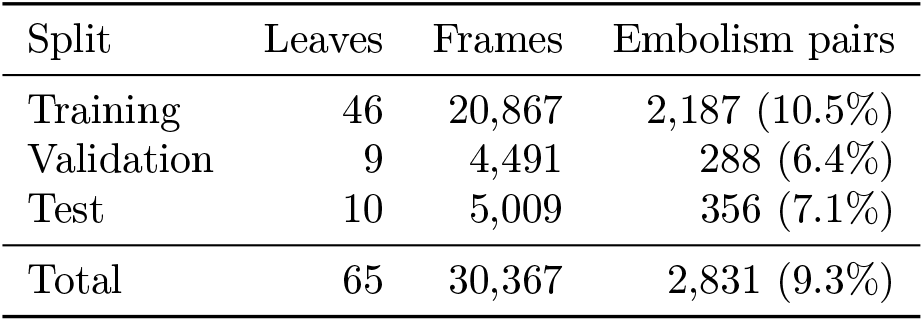
Composition of the *Senecio pterophorus* leaf dataset.

**Figure S1:**
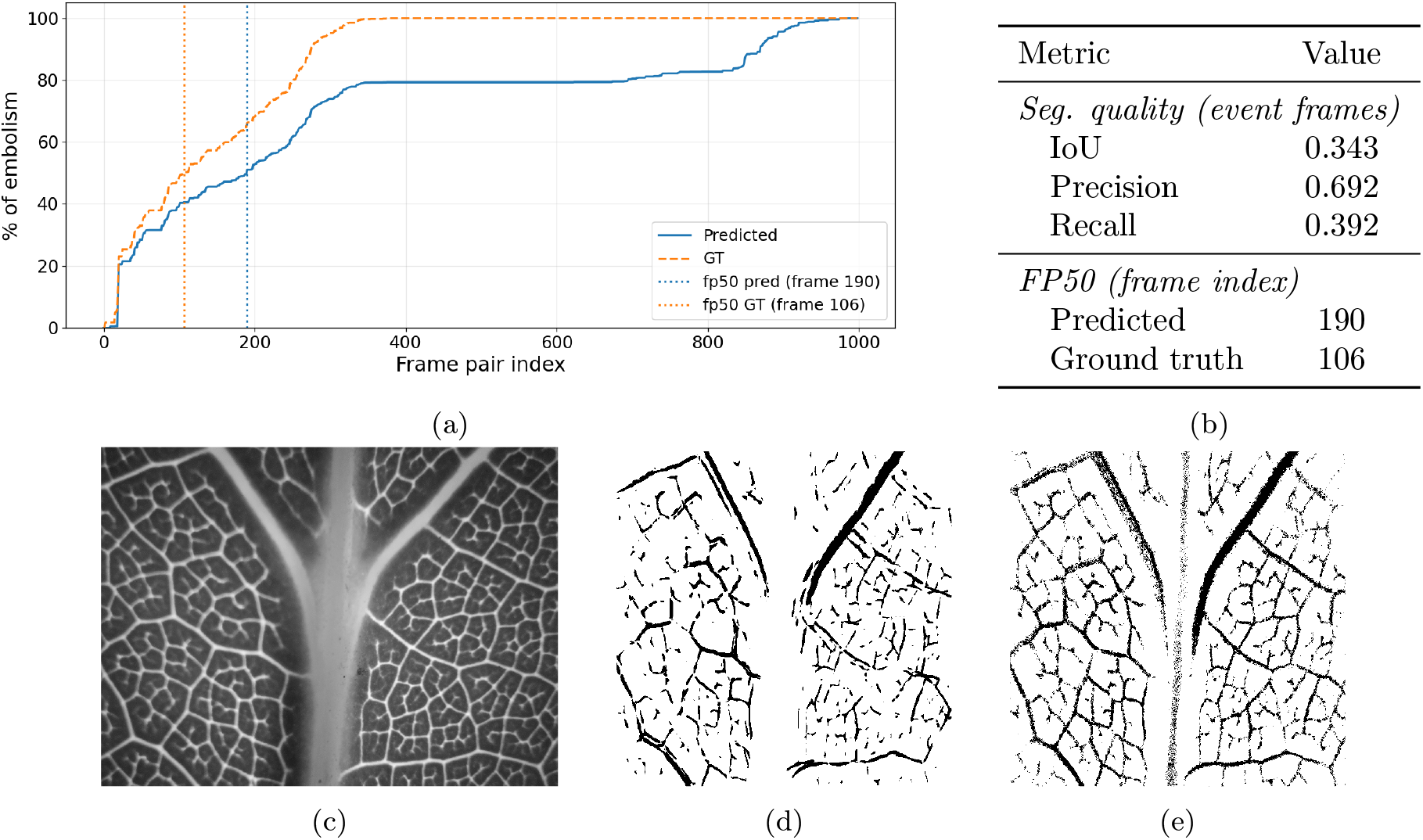
Zero-shot evaluation of the neural network model pretrained on *Senecio pterophorus* leaves and tested on leaves of an unseen species (*Liriodendron tulipifera*). **(a)** Time course of %embolism over frame indices for predicted outputs and ground truth; **(b)** pixel-level segmentation and FP50 metrics; **(c)** example of a leaf instance from the unseen species; **(d)** predicted cumulative mask; **(e)** corresponding ground truth cumulative mask.

**Figure S2:**
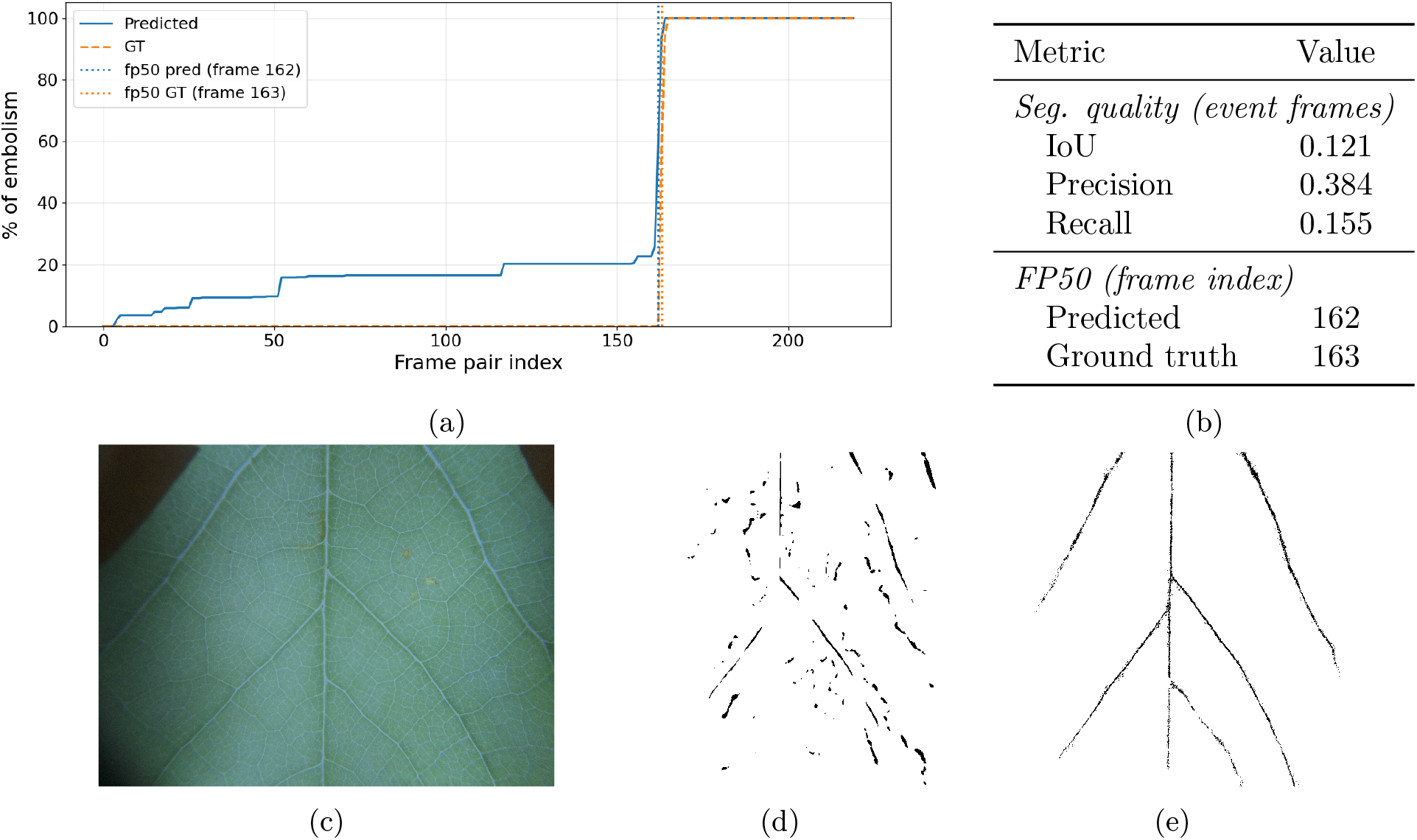
Zero-shot evaluation of the neural network model pretrained on Senecio pterophorus leaves and tested on frozen-and-thawed leaves of an unseen species (*Liriodendron tulipifera*). **(a)** Time course of %embolism over frame indices for predicted outputs and ground truth; **(b)** pixel-level segmentation and FP50 metrics, and predicted vs. ground truth FP50 metrics; **(c)** example of a leaf instance from the unseen species; **(d)** predicted cumulative mask; **(e)** corresponding ground truth cumulative mask.

**Figure S3:**
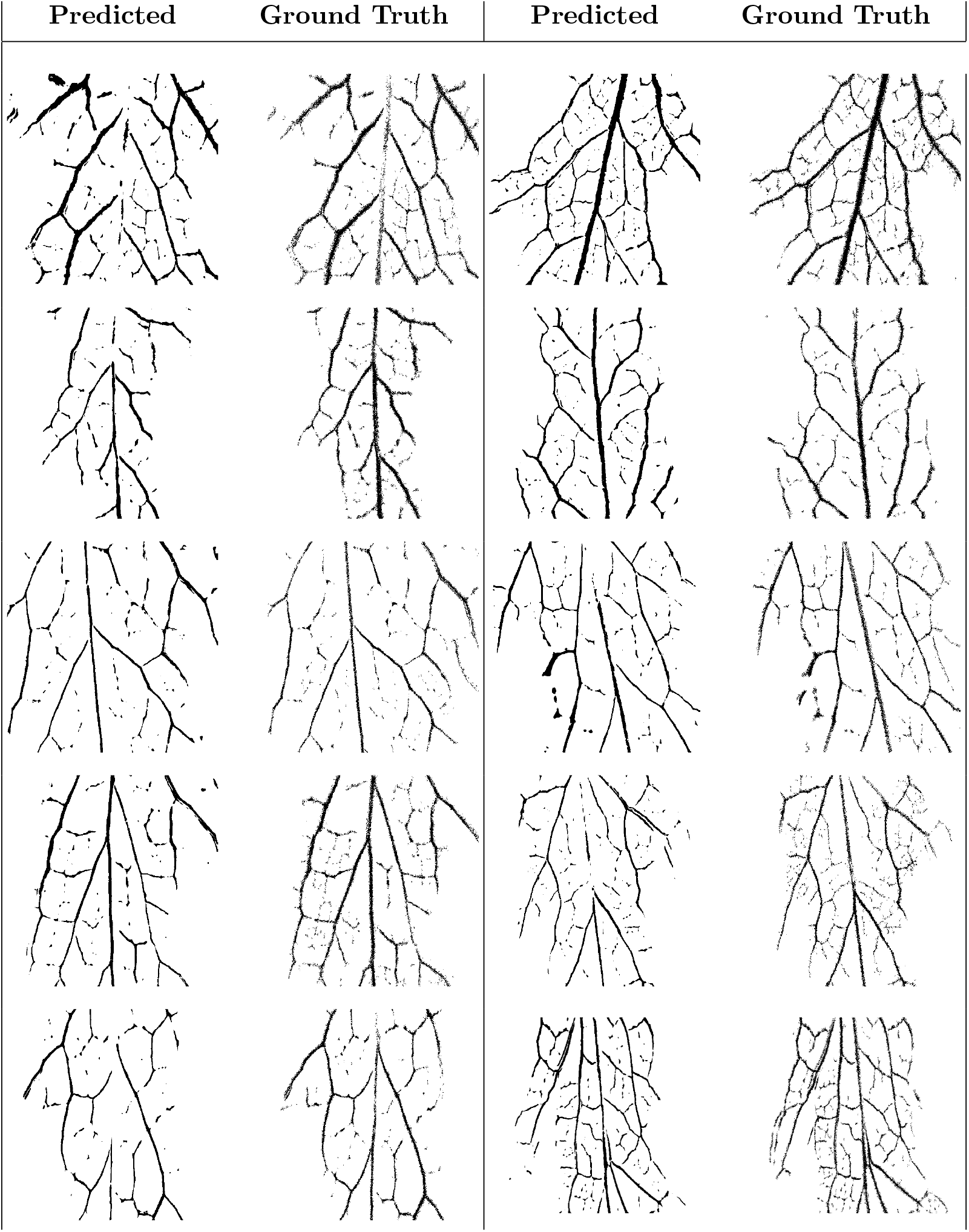
Qualitative comparison showing the difference between the model predicted cumulative embolism mask and the ground truth for 10 selected *Senecio pterophorus* leaves.

