## Supporting Information for "A data-driven approach to automate embolism detection in leaves"

#### 1 Training Details

The model is a three-level U-Net operating on a six-channel input obtained by concatenating two consecutive RGB frames. Each encoder stage applies a single $3 \times 3$ convolution with batch normalisation and ReLU followed by $2 \times 2$ max-pooling (channel progression $6 \to 32 \to 64 \to 128$, bottleneck 256). The decoder mirrors it with bilinear upsampling, skip connections, and a final $1 \times 1$ convolution with sigmoid activation, for a total of 0.97M trainable parameters. Dropout ($p = 0.3$) is applied after each decoder convolution to discourage memorisation of training-set embolism patterns.

The UNet was trained using the Adam optimiser with a learning rate of $4 \times 10^{-4}$ and a weight decay of $10^{-4}$, minimising the combined Focal–Dice loss:

$$0.5\,\mathcal{L}_{\text{focal}} + 0.5\,\mathcal{L}_{\text{dice}}$$

with $\alpha = 0.25$, $\gamma = 2.0$ and Dice smoothing 1.0.

The learning rate was halved whenever the validation loss failed to improve for five consecutive epochs, and gradients were clipped to a maximum norm of 1.0 to stabilise optimisation. Training ran for a maximum of 150 epochs with early stopping (patience 15 epochs, minimum improvement $10^{-5}$)). At inference, the predicted probability threshold is 0.5.

All experiments were run on a single NVIDIA GeForce RTX 5090 GPU. Training required approximately 2.5 hours.

### 2 Data split detail

The dataset is organised hierarchically: each leaf belongs to a mother plant (its maternal genotype), and each mother plant belongs to a source population, so that leaves from the same plant are siblings. The 65 leaves were divided into 46 training, 9 validation and 10 test sequences (Table S1). The split was not random. Beyond keeping individual leaves disjoint, the test set was deliberately built to probe the model's ability to generalise to genetic material it has never seen during training. The test sequences span three increasingly demanding levels of generalisation: five leaves come from a genotype belonging to a population that never appears in training or validation (completely unseen genetic material); four leaves correspond to a population-genotype combination absent from training; and one leaf belongs to a genotype that is represented in training by its siblings from the same plant. The validation set was constructed under the same principle, comprising one entirely unseen population and one unseen population-genotype combination whose genotype occurs in training through a different population. This graded design lets us assess not only per-leaf generalisation but also robustness to unseen populations and genotypes, and it guarantees that no population assigned to validation or test contributes any leaf to training.

Table S1: Composition of the *Senecio pterophorus* leaf dataset. *Frame pairs* are consecutive image pairs (the unit on which the model operates) and *Embolism pairs* are those whose ground-truth mask contains at least one newly embolised pixel.

| Split | Leaves | Frames | Embolism pairs |
|---|---|---|---|
| Training | 46 | 20,867 | 2,187 (10.5%) |
| Validation | 9 | 4,491 | 288 (6.4%) |
| Test | 10 | 5,009 | 356 (7.1%) |
| Total | 65 | 30,367 | 2,831 (9.3%) |

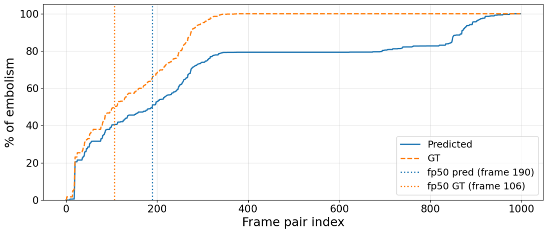


(a)

| Metric | Value |
|---|---|
| *Seg. quality (event frames)* | |
| IoU | 0.343 |
| Precision | 0.692 |
| Recall | 0.392 |
| *FP50 (frame index)* | |
| Predicted | 190 |
| Ground truth | 106 |

(b)

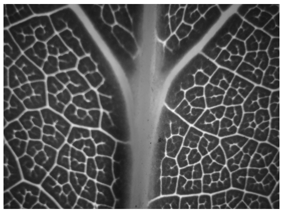

(c)

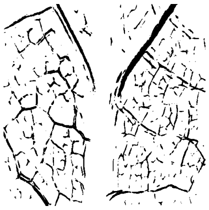

(d)

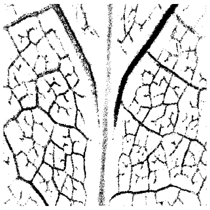

(e)

Figure S1: Zero-shot evaluation of a neural network model pretrained on Senecio leaves and tested on leaves of an unseen species (Liriodendron tulipifera). **(a)** Comparison of discrete FP-50 over frame indices for predicted outputs and ground truth; **(b)** segmentation and FP50 metrics; **(c)** example of a leaf instance from the unseen species; **(d)** predicted cumulative mask; **(e)** corresponding ground truth cumulative mask.

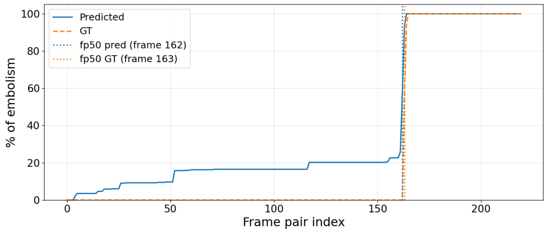


(a)

| Metric | Value |
|---|---|
| *Seg. quality (event frames)* | |
| IoU | 0.121 |
| Precision | 0.384 |
| Recall | 0.155 |
| *FP50 (frame index)* | |
| Predicted | 162 |
| Ground truth | 163 |

(b)

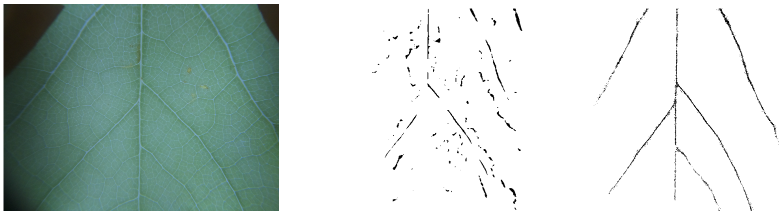

(c) (d) (e)

Figure S2: Zero-shot evaluation of a neural network model pretrained on Senecio leaves and tested on frozen-and-thawed leaves of an unseen species (Liriodendron tulipifera). **(a)** Comparison of discrete FP-50 over frame indices for predicted outputs and ground truth; **(b)** segmentation and FP50 metrics, predicted vs. ground truth; **(c)** example of a leaf instance from the unseen species; **(d)** predicted cumulative mask; **(e)** corresponding ground truth cumulative mask.

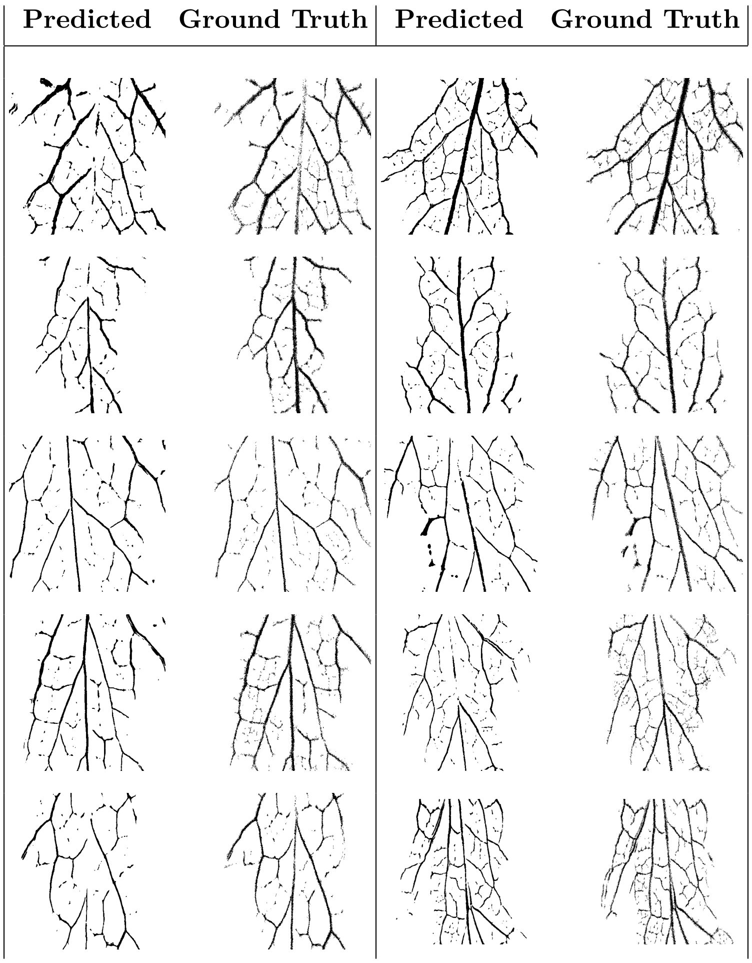


Figure S3: Qualitative comparison of the testset showing the difference between the predicted cumulative embolism mask and the ground truth (the embolism mask derived from expert image processing using the semi-automated pipeline) for 10 selected leaves from the testset dataset.
